# Multi-Scale Contextual Attention for Robust Crop and Pest Image Classification

**DOI:** 10.64898/2026.04.24.720764

**Authors:** Muhammad Majid, Hassan Tariq, Imran Mumtaz, Hanan Aljuaid

## Abstract

Image-based crop and pest recognition is considered useful for reducing the delay and cost of manual field scouting, therefore supporting timely intervention in precision-agriculture workflows. However, the real field imagery remains challenging due to the cluttered backgrounds, occlusions, illumination changes, and strong scale variation that are frequently observed across crops. The symptoms are often small or low-contrast, and pests may be partially hidden, which reduces the reliability when the setting is outside controlled environments. A unified multi-class crop–pest/condition recognition framework is presented, where a ResNet-50 backbone is utilized and enhanced with a Multi-Scale Contextual Attention (MSCA) module. The novelty is mainly considered to be achieved through the integration of explicit multi-scale contextual aggregation with lightweight joint channel and spatial attention by means of residual fusion, while the empirical evaluation was kept controlled under a fixed and reproducible protocol. A curated dataset of 21,404 field-style images covering 15 crop and pest/condition classes was compiled, and a leakage-aware fixed split with a held-out test set was adopted to support reproducibility. Augmentation was applied only to the training subset to improve robustness, although the validation data was not augmented in the same manner. On the held-out test set, balanced performance was achieved by the proposed approach, with about 0.93 accuracy and a macro-F1 score close to 0.94 being obtained, while established baselines such as EfficientNet, Vision Transformer, and attention-based CNN models were outperformed under identical evaluation settings. Controlled ablations were used to isolate the contribution of MSCA and augmentation under the same training configuration. These results indicate that lightweight multi-scale contextual attention is effective for crop and pest recognition under realistic field conditions, although some visually similar classes remained difficult.

## Introduction

Crop pests and diseases reduce crop quality and yields worldwide, yet routine field scouting is still largely manual in many regions. The process is slow and costly and early symptoms are sometimes missed, especially when weather conditions change quickly and expert support is limited for smallholder farms [1]. Scalable monitoring is therefore considered essential for meeting rising food demand. Recent advances in computer vision and deep learning have made automated visual analysis more feasible than before. When deployed on smartphones, drones, or fixed cameras, such systems can support earlier intervention, more precise pesticide use, and improved traceability across the supply chain [2].

Developments in visual recognition have been propelled by deep learning, with convolutional neural networks (CNNs) being central to them, though architecture directions have changed regularly over time. During various epochs, the models that were proposed and developed include LeNet −5, AlexNet, VGG, Inception/GoogLeNet, DenseNet, and ResNet. ResNet-50 became one of the most popular architectures due to the fact that it provided a compromise between representational and optimization stability. The residual learning method enabled the training of the network to stabilize training and refine image classification with hierarchical representations of features compared to the previous deep architectures, which had problems with optimization. The success of this architecture did not have a direct equivalent when the imagery is liberated.

In field conditions, fined grained patterns became hard to identify. Samples had cluttered backgrounds, overlapping leaves, uneven illumination and often changed scale. Discriminative cues often took up a small and irregularly placed part of the image and added confusion between classes that were visually similar and biased the model toward irrelevant parts. These challenges were also compounded by the imbalance of classes and noisy labels, which were the factors that dictated studies and which are seldom highlighted.

The problem is similar to the fine-grained classification in the background clutter and large scale difference with a computer vision point of view. Modern architectures have obtained strong results in laboratory settings, and the robustness in real settings hinged on the aggregateation of contextual information, the direction of attention mechanisms and the inhibition of the background responses. The task was decomposed by some pipelines into crop identification followed by pest or disease recognition, whereas narrow datasets were relied upon by others. When deployed in the field, robustness was reduced by such staged designs, unlike unified approaches that avoided this separation [3].

A unified multi-class classification framework was proposed to address this gap, with crop and condition patterns being jointly modeled within a single network. Better alignment with deployment constraints was achieved by this approach, and error propagation across stages was avoided, although additional optimization challenges were introduced under limited hardware and noisy supervision [4]. Augmented with a lightweight *Multi-Scale Contextual Attention (MSCA)* module was a ResNet-50 backbone, where contextual information was aggregated across multiple receptive-field scales and features were refined through joint channel–spatial attention with residual fusion. Stable optimization behavior was preserved, while diagnostically relevant regions were being emphasized through this design. Instead of introducing a new backbone family, the novelty was mainly considered in the integration of multi-scale contextual attention within a standard CNN architecture, and a controlled empirical evaluation was conducted under a fixed and reproducible protocol, with efficiency being maintained for deployment-oriented use cases.

Evaluation was being carried out on an in-house dataset of 21,404 annotated images, spanning the 15 crop and condition labels. A fixed and reproducible splitting protocol was being utilized, including the held-out test set, for preventing data leakage during training and tuning. Preprocessing was being kept consistent across all splits, and targeted augmentation was being applied only on the training subset, mainly for improving generalization, while avoiding inflating the evaluation performance in an unintended way. Although the experiments focus on agricultural imagery, the proposed attention refinement and evaluation methodology is task-agnostic and can be applied to other fine-grained image classification problems involving background clutter, and scale variation.

The main contributions of this work are summarized as follows:

- A unified 15-class crop and pest/condition recognition framework is proposed for realistic field imagery, avoiding laboratory-style constraints and staged crop–disease pipelines.
- A lightweight Multi-Scale Contextual Attention (MSCA) module is introduced, combining parallel multi-scale contextual aggregation, joint channel–spatial refinement and residual fusion within a standard ResNet-50 backbone. The design is considered to be improving feature focus under scale variation and background clutter without introducing heavy architectural complexity.
- Controlled component-level ablation experiments are conducted under a fixed and reproducible protocol. Consistent performance gains of the MSCA module over the non-attention baseline are demonstrated across multiple seeds, with statistically significant improvements, in macro-F1.
- Deployment-oriented profiling is provided, reporting parameter count, FLOPs, and inference latency alongside recognition performance to support practical decision-support use cases in precision agriculture.

The remainder of this paper has been organized as follows. In Section **??**, related work on deep learning for crop and condition recognition is reviewed. In Section **??**, the dataset, preprocessing and augmentation steps, the proposed ResNet-50 + MSCA architecture, and the training and evaluation protocols are described. Section **??** is used to report experimental results ablation studies, and comparative analyses. In Section **??**, key findings, limitations and possible directions for future work are summarized.

## Literature Review

This section is focused on image-based crop pest and disease recognition under real field conditions. Early control and yield protection are dependent on timely identification in practice. Manual diagnosis was often slow and subjective. Scaling across farms was difficult. Recent computer-vision approaches were developed to automate recognition from leaf and field imagery. Field deployment was remaining challenging, with cluttered backgrounds, varying illumination, frequent occlusion, and subtle symptoms being observed across scenes [2, 3]. Key directions in the literature were being summarized. Methodological gaps were being identified, motivating the unified multi-class formulation, and supporting the controlled study of attention mechanisms, under realistic field conditions.

### 0.1 Classification using CNN and the transfer learning Technique

Hierarchical feature learning has proven to be more successful in term capturing relevant information and thereby commencing the universal use of Convolutional Neural Networks (CNNs) in plant and crop identification when compared to hand-crafted descriptors, textures, edges, and lesion patterns. Initial studies have shown that deep models can generalise effectively in case of sufficient data and the right regularisation are in place [5]. Subsequently, the method of transfer learning had become the norm in agriculture settings, in which pre-trained backbones are fine-tuned on farm or pathology data, and frequently achieve high-quality performance even with small quantities of labelled samples. Multiple reports indicated that multi-crop and multi-disease classification tasks are highly accurate when performed using this methodology [6]. However, the loss in performance was still seen in comparative studies of AlexNet, VGG, and GoogLeNet-style designs when the symptoms were small, low-contrast, or visually ambiguous, although deeper hierarchies were used.

There was one critical limitation, however. Many pipelines often trained and evaluated on small datasets with narrow scope or control over these datasets, which overestimated real-world performance and limited generalisation across farms, seasons and imaging devices [3]. Many pipelines often trained and evaluated on small datasets that had narrow scope, or constraint, over such datasets, and this led to overestimation of the real world performance, and constrained extrapolation between farms, seasons and imaging devices [**?**]human—¿Many pipelines were also trained and evaluated on small datasets with narrow scope or control over such datasets, which overestimated real-world performance Though these limitations prompted an emphasis on diagnostically relevant cues with visual variability, this emphasis was not always obtained, and the difference between the reported accuracy and field reliability still persisted.

### 0.2 Attention mechanisms for the fine-grained recognition

Field imagery task did not only solve the assignment of a label as a class but localisation of discriminative evidence in the scene as well. The distractors often included soil texture, shadows and the vegetation at the back of the models leading to the drift of attention to irrelevant parts. As a control to this effect, attention mechanisms were added to re-weight features and focus on informative regions including lesion clusters, pest presence, and discoloration patterns. The formulations of residual attention were incorporated in CNN backbones, where the feature refinement did not destabilise the optimisation. As a result, the response to minor symptoms was more sensitive, but the rejection of irrelevant texture was not always similar in all cases.

Multi-scale processing was considered critical for fine-grained recognition. Target size and visibility were varied. The same condition was observed to appear differently across growth stages. Pyramid-style designs with adaptive feature fusion were explored to capture small or crowded targets across scales [7]. Related fusion modules were proposed for dense prediction tasks. For example TBSFF-style mechanisms were designed to combine multi-scale context with fine detail to improve segmentation quality [8]. In recognition settings discriminative feature responses were emphasized by channel attention. Background noise was reduced and localization of symptom patterns was improved by spatial or coordinate attention. A SqueezeNext variant using multi-scale kernels with coordinate attention was reported to produce fewer false positives and sharper delineation of affected regions [9]. Collectively attention mechanisms were shown to improve feature focus and robustness to scale variation. However, the individual contribution of these components was often difficult to assess as the datasets splits and training protocols were varied across studies.

### 0.3 Edge friendly models and the evaluation benchmarks

Limited compute resources, unstable power availability, and constrained memory capacity often restricted practical use in real field deployments. To cope with these constraints, lightweight and efficiency-focused designs were explored, aiming to reduce inference time while maintaining accuracy. For real-time inference on embedded platforms, mobile-oriented backbones—particularly MobileNet-style models—were therefore adopted [10]. Through pruning, quantization, and knowledge distillation, additional efficiency gains were achieved, enabling deployment on drones, tractors, and greenhouse cameras, although the resulting trade-offs were not always fully accounted for.

Steadily, progress in this area continued, shaped by stronger datasets and increasingly rigorous evaluation protocols, even if direct translation to real-world reliability did not always occur. For broader comparison and protocol standardization, large-scale benchmarks such as the AI Challenger 2018 Crop Disease Detection dataset were utilized. Using a dual-branch model with efficient channel attention, competitive benchmark performance was reported [11]. Label quality and dataset splitting strategies were revisited in later studies, where external images were incorporated and lightweight training schemes were adopted to improve balance and generalization [12]. Greater emphasis was placed on leakage-free splits, cross-domain evaluation (e.g., cross-farm or cross-season testing), and module-level ablations to isolate architectural effects. Still, key challenges remained: long-tail imbalance persisted, small or densely clustered targets were difficult to recognize, and the trade-off between accuracy and real-time constraints continued as a practical limitation [13].

### 0.4 Summary of gaps and motivation

As observed in literature, two gaps are repeatedly observed in practical field settings. First, controlled comparisons under identical data splits and training protocols are often not reported. Because of this, performance gains are sometimes attributed to architectural components even when the protocol was different. Second, deployment-oriented claims are frequently made without reporting model size or parameter count or inference latency. As a result, reproducibility is claimed while practical justification is not fully supported.

A unified multi-class formulation is emphasized in the present work. A lightweight multi-scale contextual attention refinement is also emphasized. A controlled empirical evaluation is conducted to isolate the effects of attention design and data augmentation. However, these effects are isolated under a fixed and reproducible protocol, which may still vary across experiments.

## Methodology

This section describes the pipeline that was utilized for building and evaluating the proposed crop–condition recognition system. A curated dataset of 21,404 field-style images spanning 15 labels was assembled and was quality-checked during preprocessing. Reproducibility was treated as a first-class requirement while designing the workflow. For that reason, the dataset was split into training, validation, and a held-out test set before any augmentation, using a fixed random seed and stratified sampling [14]. At the outset, before any training was carried out, resizing and normalization of images were performed. Only on the training split was augmentation applied, with field variability being emulated, while evaluation performance was not being inflated.

In the core design, a ResNet-50 feature extractor was being used, augmented with the Multi-Scale Contextual Attention (MSCA) module, and followed by a lightweight classification head, producing 15-way predictions. Based on validation performance, hyperparameters were selected. Once on the frozen test set, final performance was reported, using accuracy, macro-precision, macro-recall, macro-F1, and Matthews Correlation Coefficient (MCC) [15]. In Figure 1, the end-to-end workflow was summarized, while the overall processing stages were being illustrated.

**Fig 1.**
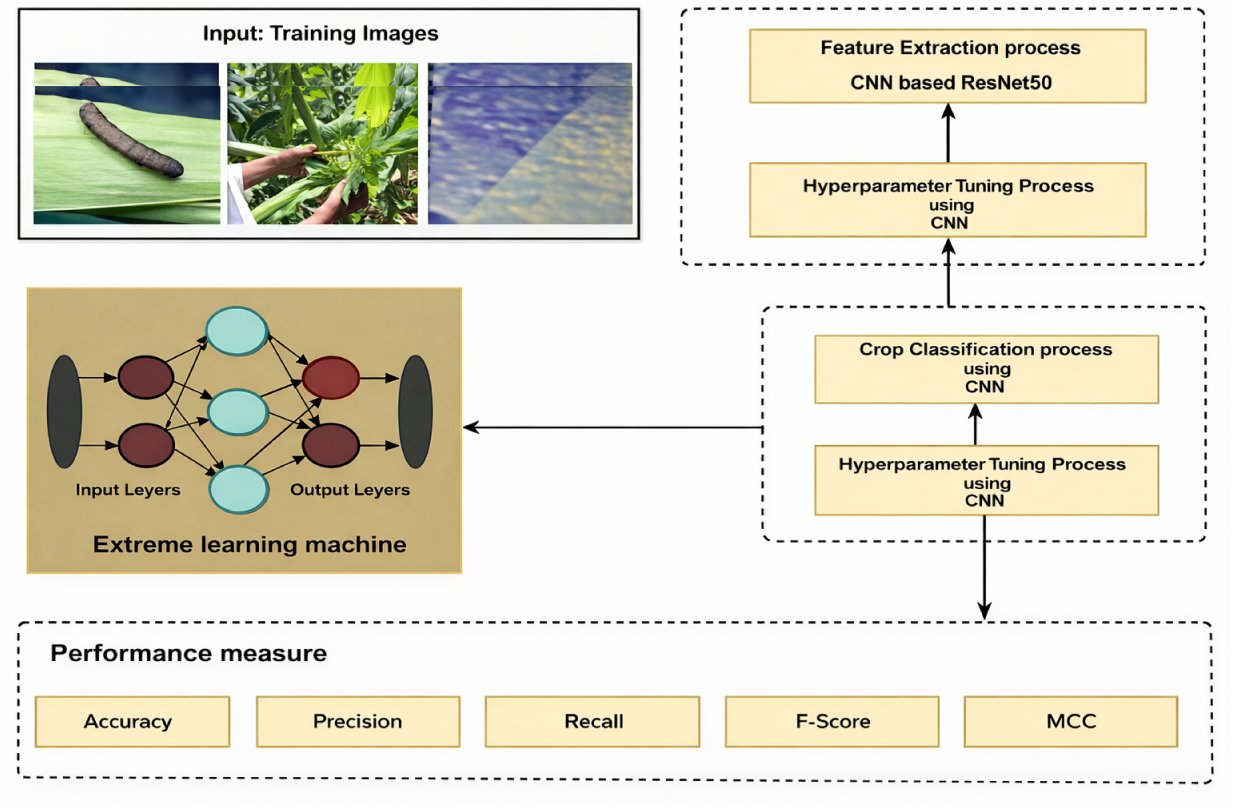
Overview of the proposed pipeline: dataset curation and split, training-only augmentation, ResNet-50 + MSCA feature learning, and 15-class prediction with reproducible evaluation.

### 0.5 The Data acquisition, labeling, and the split protocol

#### 0.5.1 Dataset source and final 15-label taxonomy

Images in this study have been chosen based on the dataset that has been cited in the article of [16] that contains the images of agricultural fields under natural conditions. The acquisition was done on handheld devices as well as field cameras.

In the gathered images, there was observed diversity in terms of illumination, background clutter hence real deployment conditions instead of controlled laboratory environments. As a result, there were differences in the viewing angles and the visibility of symptoms that were occasionally very high.

Based on this source, a single 15-label taxonomy was built with various crops and pest or disease states. To reduce ambiguity and balance in the classes, visually similar and low-frequency classes were combined; however, there were still some controversial points of label boundaries. Table 15 labels were administered consistently and its spread is reported in Table 1.

**Table 1.** Final 15-label taxonomy and class distribution used in this study (stratified 70–15–15 split).

| ID | Label name | Total | Train | Val | Test |
| --- | --- | --- | --- | --- | --- |
| 0 | Corn Common Rust | 1,400 | 980 | 210 | 210 |
| 1 | Corn Gray Leaf Spot | 1,300 | 910 | 195 | 195 |
| 2 | Healthy Corn | 2,000 | 1,400 | 300 | 300 |
| 3 | Corn Pest (Aphids/others) | 1,500 | 1,050 | 225 | 225 |
| 4 | Corn Leaf Blight | 1,300 | 910 | 195 | 195 |
| 5 | Maize Ear Rot | 900 | 630 | 135 | 135 |
| 6 | Maize Stem Borer | 900 | 630 | 135 | 135 |
| 7 | Healthy Maize | 1,200 | 840 | 180 | 180 |
| 8 | Rice Pest/Condition | 2,800 | 1,960 | 420 | 420 |
| 9 | Wheat Healthy | 1,500 | 1,050 | 225 | 225 |
| 10 | Wheat Pest/Condition | 1,400 | 980 | 210 | 210 |
| 11 | Wheat Yellow Rust | 1,400 | 980 | 210 | 210 |
| 12 | Cotton Pest/Condition | 1,400 | 980 | 210 | 210 |
| 13 | Sugarcane Pest/Condition | 1,300 | 910 | 195 | 195 |
| 14 | Millet Pest/Condition | 1,104 | 773 | 166 | 165 |
| - | **Total** | **21,404** | **14,983** | **3,211** | **3,210** |

Each image was given only one image-level label which was the dominant crop and condition. Ambiguous and poor quality samples were eliminated, thus improving label reliability, but maintaining field variability. Overall, the dataset included 21,404 images, and when partitioning the 70-15-15 splits were considered as target proportions not as constraints.

Representative examples for the 15 labels are shown in Figure 2. The figure illustrates visual diversity, symptom subtlety, and background complexity under field imagery.

**Fig 2.**
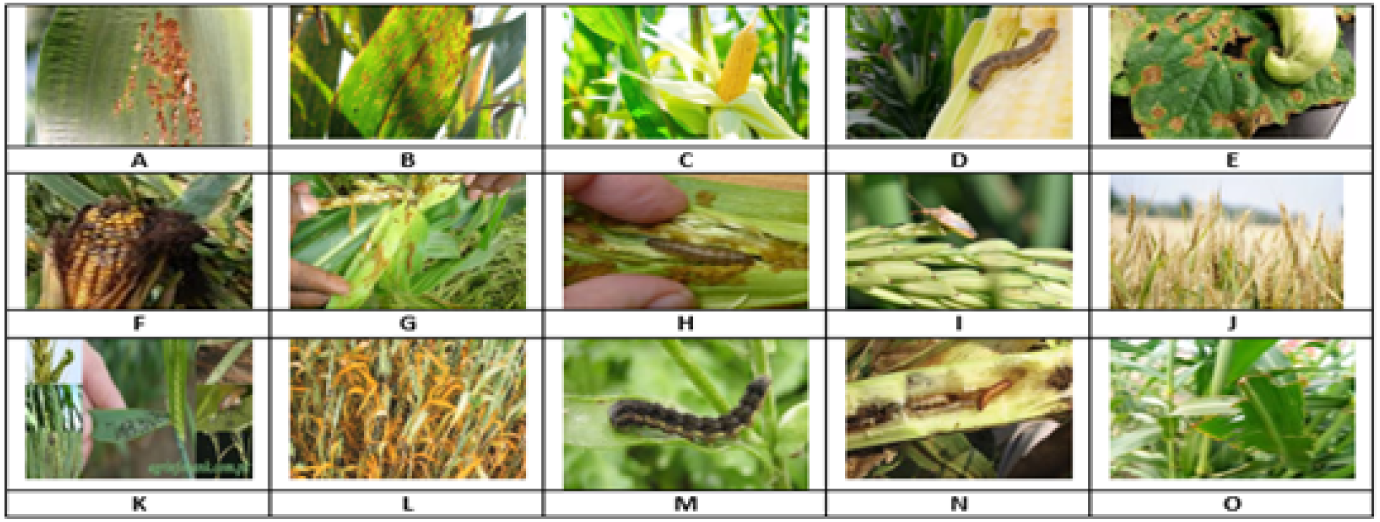
Representative examples for the final 15 labels used in this study (label names and counts are defined in Table 1).

#### 0.5.2 Labeling procedure for image-level classification

This work considered image-level multi-class classification, where-by the label of one element in Table 1 to each image, so that the learning task was single-label classification, as opposed to multi-instances interpretation.

The problem was not stated as object detection, and bounding boxes were not used nor predicted by the model in the training process. This explanation was added to avoid the possibility of confusing the detection style annotation tools and the desired classification goal, but the visual look of the annotation interface might still give a hint of a detection rig to the reader, especially when the experimental setup is being surveyed in a hurry.

#### 0.5.3 The Splitting protocol and leakage prevention

Through the experimental design, leakage prevention was addressed, as dataset splitting was performed before any augmentation to enforce early separation. A 70–15–15 stratified protocol was applied to preserve per-class proportions [14], and reproducibility was ensured by using a fixed random seed. Using perceptual similarity checks, near-duplicate screening was carried out; duplicates were removed or confined to a single split, although a few borderline cases still remain debatable.

Only the training set underwent augmentation. Apart from resizing and normalization, which were applied uniformly, validation and test sets were kept unchanged and were not treated as a source of leakage. Class imbalance was handled mainly through training-only augmentation, and when helpful, class weighting in the loss [17]. The test set was frozen for all experiments. Model selection relied only on validation performance.

For transparency, the file lists for training, validation, and test were actively saved as CSV files containing image paths and label IDs. Where permitted, the split CSV files and the training configuration, such as random seeds and key hyperparameters, will be released to support reproducibility. Redistribution of the raw images may be restricted due to licensing constraints of the original source dataset, but the split CSV files, label taxonomy, and training configuration will be made available to enable reproducibility and fair comparison.

### 0.6 Preprocessing and the augmentation

All images were resized to 256 × 256 RGB, and were normalized using the ImageNet channel-wise mean and standard deviation [18, 19]. The same preprocessing was being applied during training and during inference, thereby keeping processing behavior consistent. By maintaining this consistency across splits, the input distribution was being kept stable, therefore reducing the likelihood of evaluation being affected by preprocessing drift or unintended variation.

#### 0.6.1 Training-only augmentation (with explicit ranges)

Field images were observed to vary strongly in pose, scale, and lighting. Augmentation was therefore utilized to improve robustness, and was applied only on the training split. Augmentation was generated on-the-fly, so that the underlying split sizes remained exactly as reported in Table 1. The configuration was summarized in Table 2, with ranges being selected to mimic plausible field variation, while avoiding unrealistic distortions, although some rotations were still considered slightly unnatural in certain crop layouts.

**Table 2.** Training-time augmentation configuration (applied only on the training split).

| Augmentation | Setting / range |
| --- | --- |
| Random rotation | $\pm 25^\circ$ |
| Random zoom | [0.90, 1.10] |
| Width/height shift | $\pm 10\%$ of image size |
| Horizontal flip | Enabled (probability 0.5) |
| Brightness jitter | $\pm 15\%$ |
| Contrast jitter | $\pm 15\%$ |
| Gaussian noise | $\sigma \in [0, 0.02]$ (applied with prob. 0.3) |

The split-before-augment workflow is visualized in the given Figure 3. While the representative augmented samples are shown to illustrate the type of variability presented during training. However, the augmented examples are not intended to represent the full range.

**Fig 3.**
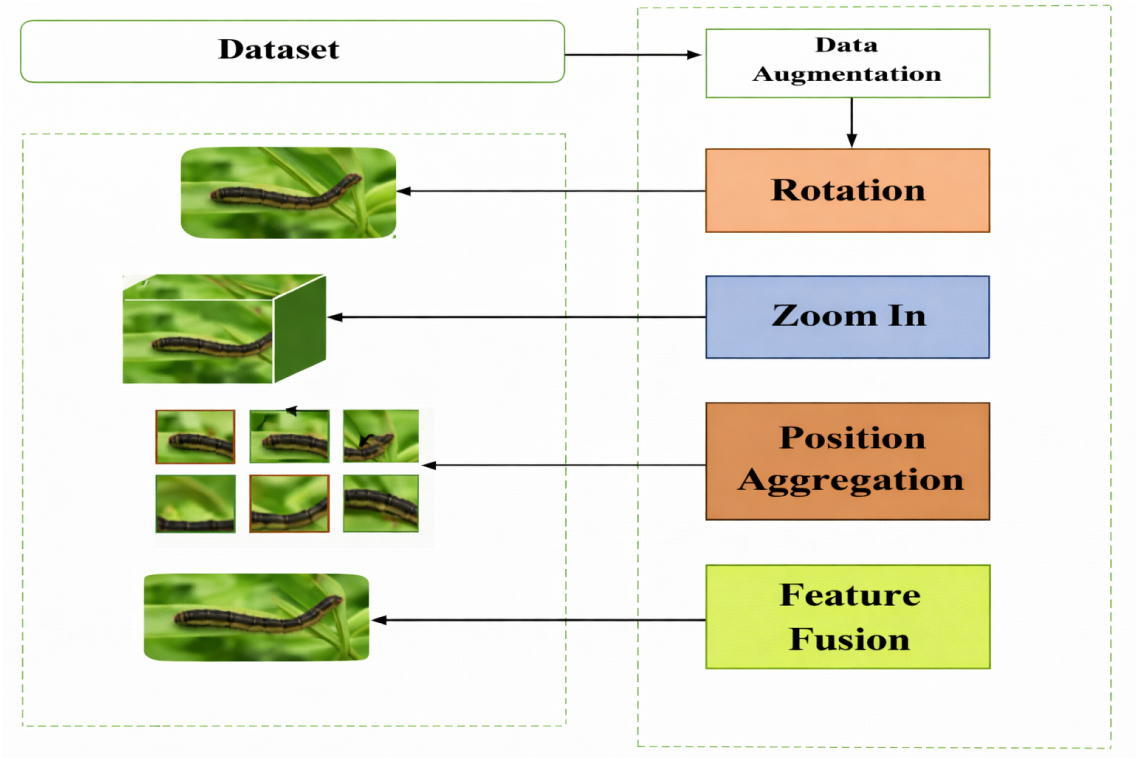
Dataset preparation: split-before-augment to prevent leakage; augmentation applied only to training images; validation/test kept unchanged except resize/normalization.

#### 0.6.2 Optional image-mixing augmentation (feature fusion)

In addition to standard transforms, optional image-mixing was evaluated as a regularizer. It was enabled only when validation macro-F1 improved. Unless stated otherwise, the primary results correspond to the standard augmentation setting, and image-mixing is treated as an ablation variant.

Let (*x_i_, y_i_*) and (*x_j_, y_j_*) be two training samples with one-hot labels. A MixUp-style fusion is defined as:

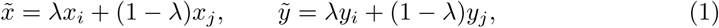

where *λ* ∼ Beta(*α, α*) with *α* = 0.2 in our implementation. For a CutMix-style variant, a rectangular region is replaced and *λ* is proportional to the retained area. These fusions are applied only within the training split [20, 21]. They are never applied to validation or test data.

### 0.7 Model architecture

The proposed architecture contains three components. First is a ResNet-50 backbone. Second is the MSCA attention module. Third is a lightweight classification head that produces logits for 15 classes. The ordering is simple, although the attention block is placed inside the backbone feature stream.

#### 0.7.1 Backbone: ResNet-50 feature extractor

ResNet-50 was adopted to extract discriminative features while keeping computational cost moderate [22]. ImageNet-pretrained weights were used for transferring generic visual structure, and the network was then fine-tuned for adapting to field conditions such as clutter illumination shifts, and partial occlusion [23]. Inputs were resized to 256 × 256 and were normalized using ImageNet statistics [18, 19].

The canonical ResNet-50 topology was retained throughout the experiments. A 7 × 7 convolution with stride 2, followed by a 3 × 3 max-pooling operation, was used for forming the stem [24]. Four residual stages were then used for producing progressively lower-resolution feature maps. Intermediate features at 16 × 16 resolution were forwarded to the MSCA module, which was considered effective for preserving spatial detail. This stage was selected as providing a practical balance between spatial resolution and semantic richness, whereas earlier layers were considered too low-level, and later layers were observed to lose fine details due to repeated downsampling. Final 8 × 8 features were globally pooled for obtaining a 2048-dimensional representation [25]. In a few preliminary runs, features from earlier stages were considered, but were not retained due to unstable behavior being observed.

Fine-tuning was performed in two stages. First, the backbone was frozen and only the head was trained. Second, upper stages were unfrozen gradually with a smaller learning rate than the head. Weight decay and early stopping were used. Checkpointing was driven by validation macro-F1. Mixed precision was enabled when supported to improve throughput and reduce memory usage [26]. All settings affecting reproducibility were logged, including seeds, unfreeze schedule, and learning rates [27].

#### 0.7.2 Attention: Multi-Scale Contextual Attention (MSCA)

MSCA is designed to improve symptom-focused learning under scale variation and clutter. Diagnostic cues are emphasized while stable optimization is preserved through residual fusion [28]. While the components are related to established multi-scale and attention mechanisms, the novelty here lies in their lightweight integration within a standard CNN and in the controlled evaluation that isolates their contribution under a fixed protocol. To further clarify the novelty of the proposed MSCA module, it is compared with attention mechanisms such as Squeeze-and-Excitation (SE), CBAM, and pyramid attention designs.

The SE module focuses on channel recalibration. The spatial structure of the feature map is not explicitly considered in SE, and contextual aggregation is obtained only through global pooling. This improves channel selectivity but scale variation is not directly addressed in field imagery.

The CBAM module extends SE by incorporating spatial attention after channel attention. However, the attention operations in CBAM are performed on features extracted at a single receptive-field scale. The contextual information was therefore being limited to the inherent scale of the backbone feature map. In complex agricultural scenes, where lesions and pests were appearing at different sizes and densities, such single-scale processing was often being insufficient, even when deeper backbones were being used.

Pyramid attention mechanisms were being designed for capturing multi-scale information. However, reliance was often placed on multi-stage feature pyramids, or on heavy fusion across hierarchical levels. Such strategies were increasing model complexity and computational cost, which were being considered unsuitable for deployment-oriented agricultural systems, especially under memory constraints.

The proposed MSCA module was differentiated through three aspects. First, explicit multi-scale contextual aggregation was being performed using parallel dilated convolutions at a single intermediate feature level, with contextual information at different receptive-field sizes being obtained simultaneously and fused before attention refinement. Second, joint channel and spatial attention was being applied on the fused representation, rather than being applied sequentially. Third, refined features were being integrated with the original backbone features through residual fusion, with the original signal being preserved and stable optimization being supported during fine-tuning.

The novelty of MSCA was therefore being defined by the efficient combination of parallel multi-scale context extraction, lightweight joint attention refinement, and residual integration within a standard ResNet backbone. The objective was not framed as introducing a new backbone family, but as providing a deployment-aware refinement mechanism, with its contribution being isolated through controlled ablation. A concise conceptual comparison with SE, CBAM, and pyramid attention mechanisms was being provided in Table 3.

**Table 3.** Conceptual comparison of MSCA with common attention mechanisms.

| Method | Multi-scale Context | Channel Attention | Spatial Attention |
| --- | --- | --- | --- |
| SE | No | Yes | No |
| CBAM | No (single scale) | Yes | Yes (sequential) |
| Pyramid Attention | Yes (multi-stage) | Yes | Yes |
| Proposed MSCA | Yes (parallel dilated) | Yes | Yes (joint + residual) |

Figure 4 illustrates the MSCA design. Parallel 3 × 3 convolutions with different dilation rates capture contextual evidence at complementary receptive-field sizes. This is useful in field imagery where lesions and pests can be small, sparse, or partially occluded. The fused multi-scale features are then refined through lightweight channel and spatial attention. Background responses are suppressed and diagnostically relevant regions are emphasized. Finally, residual fusion preserves the original feature signal and supports stable gradient flow during fine-tuning.

**Fig 4.**
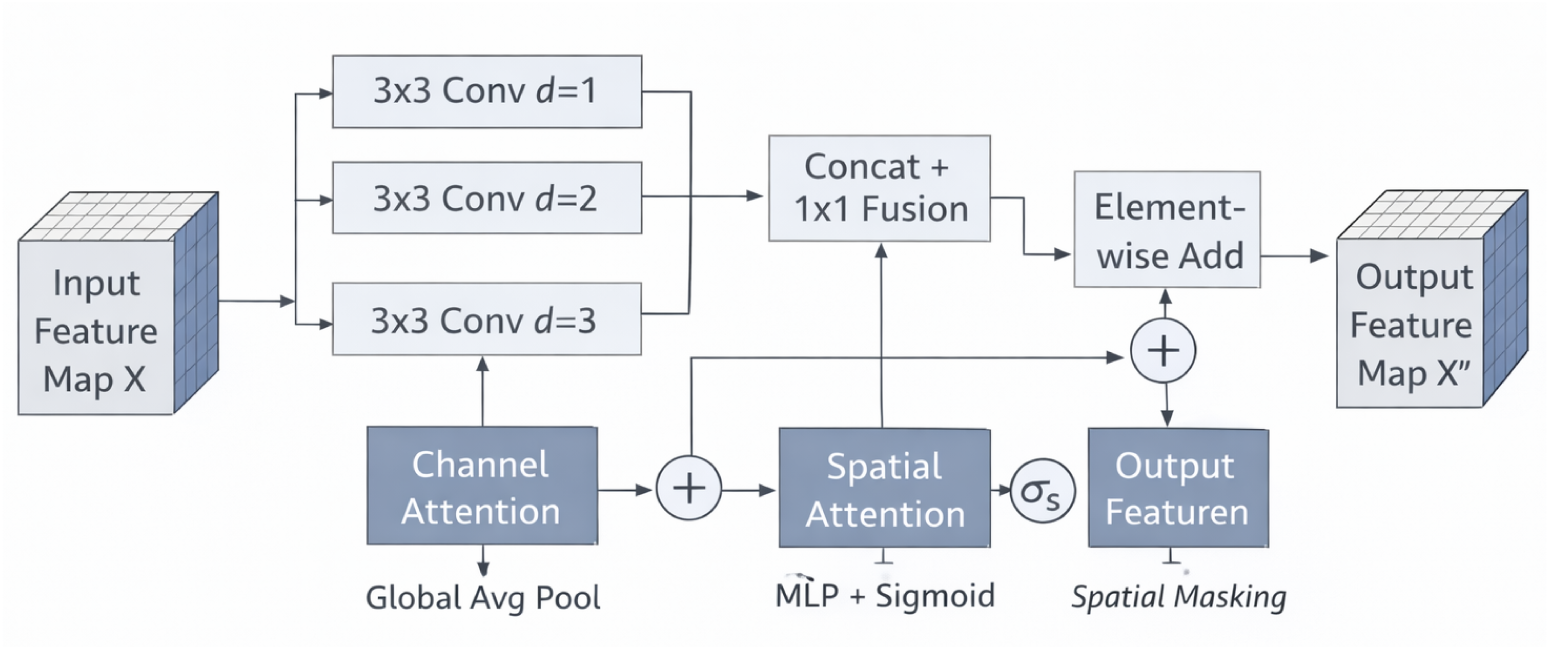
Structure of the proposed Multi-Scale Contextual Attention (MSCA) module. Parallel dilated convolutions aggregate contextual information at multiple receptive-field scales, followed by channel and spatial attention refinement. The refined representation is fused with the input feature map through residual addition to preserve stable optimization.

Let *X* ∈ ℝ*^H×W^ ^×C^* be an intermediate backbone feature map. MSCA applies *K* parallel dilated convolutions (here *K* = 3 and *d_k_* ∈ {1, 2, 3}) and fuses them:

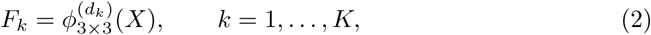

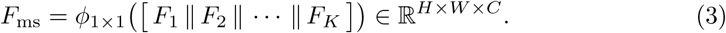

Channel attention is computed by global average pooling followed by a lightweight MLP:

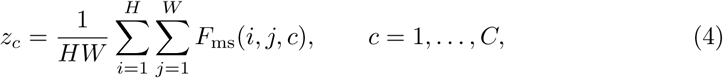

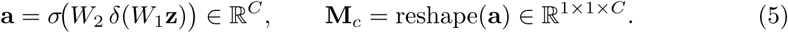

Spatial attention is obtained by pooling across channels and applying a small convolution (we use *k* = 7):

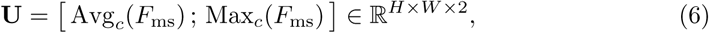

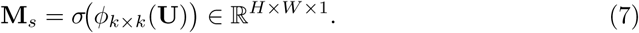

The refined output is produced via residual fusion:

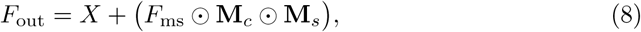

Where 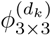 is a 3 × 3 convolution with dilation *d_k_*, *ϕ*_1*×*1_ is a 1 × 1 fusion convolution, [·∥·] denotes channel concatenation, *δ*(·) is ReLU, *σ*(·) is sigmoid, and ⊙ denotes element-wise multiplication with broadcasting.

### 0.8 Classification head, activation functions, and output

Refined features are mapped to class scores by a lightweight head. Global average pooling is applied to the final tensor *F* ∈ ℝ^8*×*8*×*2048^ to obtain *h* = GAP(*F*) ∈ ℝ^2048^. A fully connected layer produces logits *z* ∈ ℝ*^C^* for *C* = 15 classes:

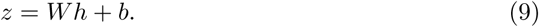

Softmax converts logits to probabilities:

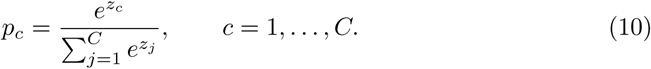

Training minimizes multi-class cross-entropy, optionally with class weights:

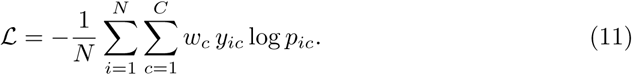

When an intermediate nonlinearity is used in the head, LeakyReLU is applied to support gradient flow under imbalance. Dropout (*p* = 0.3) is enabled only when validation macro-F1 improves. The head variants evaluated in ablations are summarized in Figure 5.

**Fig 5.**
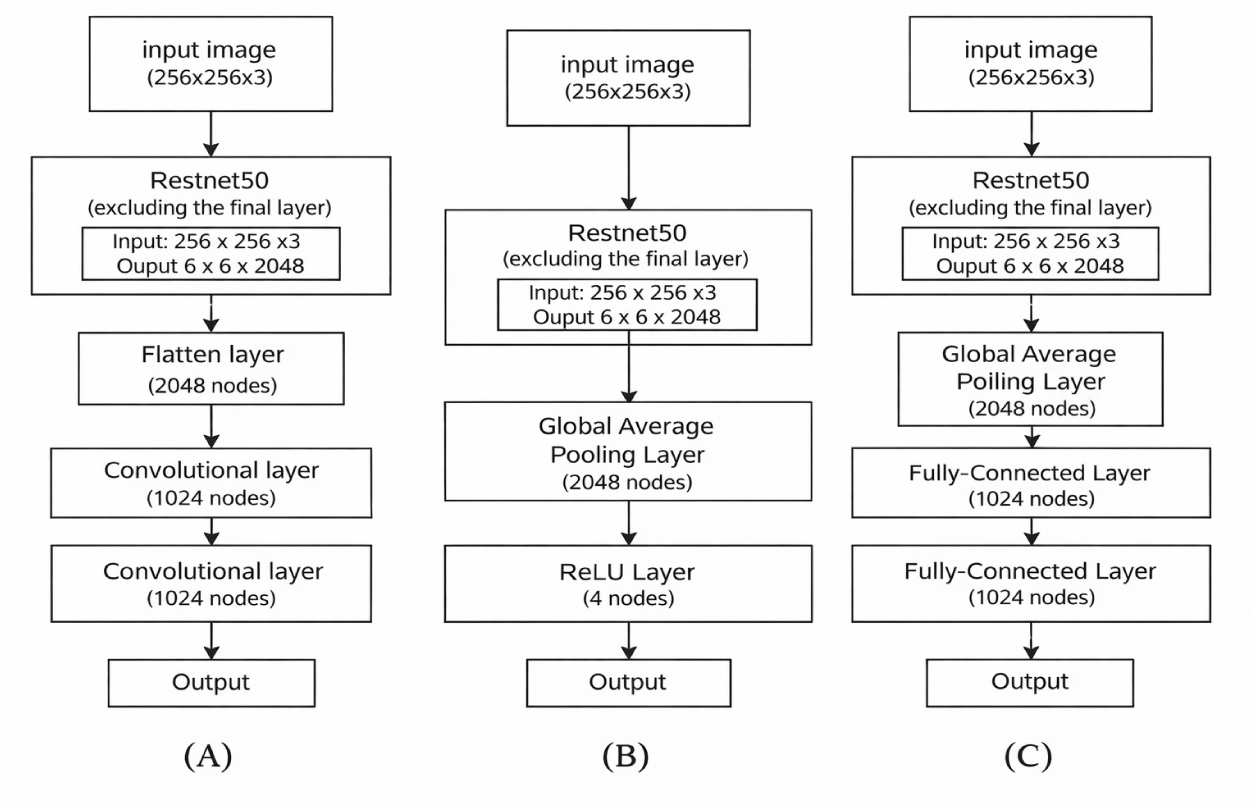
Classifier head variants evaluated: (A) Flatten head, (B) Global Average Pooling (GAP) with nonlinearity, and (C) GAP with an additional fully connected layer.

### 0.9 Hyperparameter tuning and implementation details

Hyperparameters were tuned using the validation split to improve generalization and reduce overfitting [29]. Adam was used with *β*_1_ = 0.9, *β*_2_ = 0.999, *ɛ* = 10*^−^*^8^, initial learning rate 10*^−^*^4^, and batch size 32. A cosine decay schedule was adopted. Early stopping was driven by validation macro-F1 [30]. During fine-tuning, differential learning rates were applied, with the head learning faster than the backbone.Regularization was provided through weight decay. Where supported, mixed precision was enabled to improve throughput and training stability [26]. In several runs, early stopping was triggered sooner than expected because validation behavior was noisy, meaning that the best checkpoint was not always found near the final epoch, which made comparisons across runs less straightforward.

Using Python 3.9, experiments were conducted with Keras built on top of TensorFlow and executed in Ubuntu-based environments. Training epochs were run on NVIDIA A100 and V100 GPUs, while random seeds, split CSV files, optimizer settings, and scheduler parameters were recorded to support reproducibility [31]. Summarized in Table 4 is the overall training and evaluation setup, although some configuration details were reused across experiments rather than varied.

**Table 4.** Hardware and software configuration used for training and evaluation.

| Component | Specification |
| --- | --- |
| Processor | Intel Core i7-7800 CPU |
| RAM | 16 GB |
| GPU | NVIDIA A100 / NVIDIA V100 (DGX) |
| Operating System | Ubuntu 16.04 |
| Programming Language | Python 3.9 |
| Framework | Keras + TensorFlow |
| Optimizer | Adam ($\beta_1=0.9$, $\beta_2=0.999$, $\epsilon=10^{-8}$) |
| Initial Learning Rate | $10^{-4}$ (cosine decay) |
| Batch Size | 32 |
| Head activation (if used) | LeakyReLU |
| Early stopping criterion | Validation macro-F1 |

### 0.10 Inference and deployment-oriented profiling

At the inference time every input image was resized to a square size of 256 × 256, transformed into the RGB colour space, and normalised using the same statistical parameters used in the training. A ResNet-50 backbone, Multi-Scale Contextual Attention (MSCA) module, global average pooling layer and a classification head were then applied to the pre-processed tensor, producing a probability distribution over 15 classes. In order to improve throughput in a real-world situation, a batch inference on a graphics card was conducted. Unless indicated in an ablation study, test-time augmentation was not used at all [32]. Uninterrupted preprocessing is required when deployed; inconsistencies in the required pipeline would modify the reported performance measures.

To facilitate the arguments about the suitability of deployment, the computational characteristics were profiled with the hardware configuration in Table 4. The overall number of parameters was taken in the model summary, and latency was computed as the median execution time in several forward passes after a warm-up period without data-loading delays. Standard profiling tools estimated the floating-point operations (FLOPS) of variants of the model with a fixed input resolution of (256 × 256). To control comparability, measurements were made on the same software stack and preprocessing pipeline. The given accuracies were also provided with the latency and FLOP numbers, thus, correlating recognition performance with the deployability, but variability over run time and hardware environment was still and could not be controlled completely.

### 0.11 Evaluation protocol and metrics

The evaluation had a strict protocol. After picking the model that performs best on the validation split, the model was frozen and then tested on the held-out test set a single time only [15]. No augmentation was done to the test set and normalisation and resizing were done only. Reported were accuracy, macro-precision, macro-recall, macro-F1 and Matthews correlation coefficient (MCC). Per-class metrics were averaged to obtain macro scores, and multi-class confusion Table was computed to obtain MCC. All ablation experiments and controlled comparisons were done using the same data split and random seed, with variations of the auxiliary head, augmentation strategies, the presence or absence of MSCA activation, and changes in backbone depth. This method enabled the direct comparison and reduced the possibilities of variability analysis.

In the experiments of ablation, only one component (e.g., the MSCA module) was altered, but the data split, optimiser settings, learning schedule, and random seed were kept fixed. This design allowed architectural effects to contend with performance variations, but a single frozen test set exposed sampling effects to bias; it was also vulnerable to sampling noise effects due to the reliance on only one set of tests at a time, as opposed to experimentation with multiple sets of tests at once that could be reused and pooled together later in the analysis [33].

## Results and Discussions

The proposed universal crop condition recognition system was tested on the real field images. A ResNet- 50 backbone was used with the MSCA module in the inference pipeline and it was trained on 21404 images of fifteen semantic categories. There was no control of visual conditions, cluttered backgrounds, partial occlusions and fluctuating illumination in most samples; a number of classes were harder to identify due to tiny or low-contrast manifestations of symptoms.

A held-out test split was fixed and not observed during training and hyperparameter optimization and only the last checkpoint was chosen based on validation macro-F1.

Despite the fact that more robust statistical approximations might have been achieved through cross-validation or repeated random splits, they were considered less representative of the deployment situations, in which a model is trained once and tested once using an unknown set.

Robustness was tested by stable dynamics of learning, and controlled ablation studies which performed under the same split and random seed. The comparative framework therefore enabled analysis of model variants whilst maintaining a realistic deployment scenario. Among the reported results are learning dynamics, held-out performance, MSCA and augmentation ablations, class-wise behaviour on a balanced diagnostic subset, backbone-depth comparisons, deployment-oriented profiling and an indicative comparison to recent literature, but not all these aspects were given equal weighting.

### 0.12 Learning curves and training stability

Before test performance was discussed, convergence and generalization were verified by examining the learning curves. For field imagery, this step was considered important, since deep models can learn shortcut cues from background texture when regularization is weak. Shown in Figures 6 and 7 are training and validation accuracy and loss over 20 epochs for ResNet-50 + MSCA.

**Fig 6.**
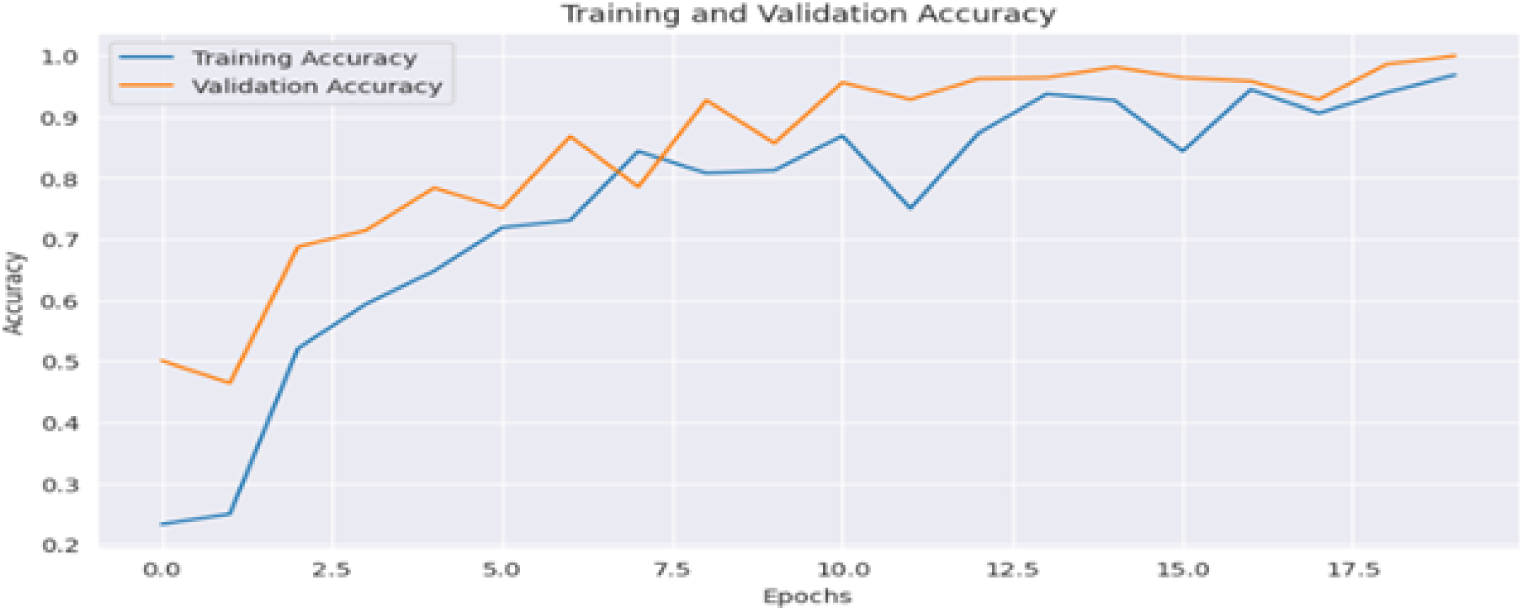
Training and validation accuracy over 20 epochs.

**Fig 7.**
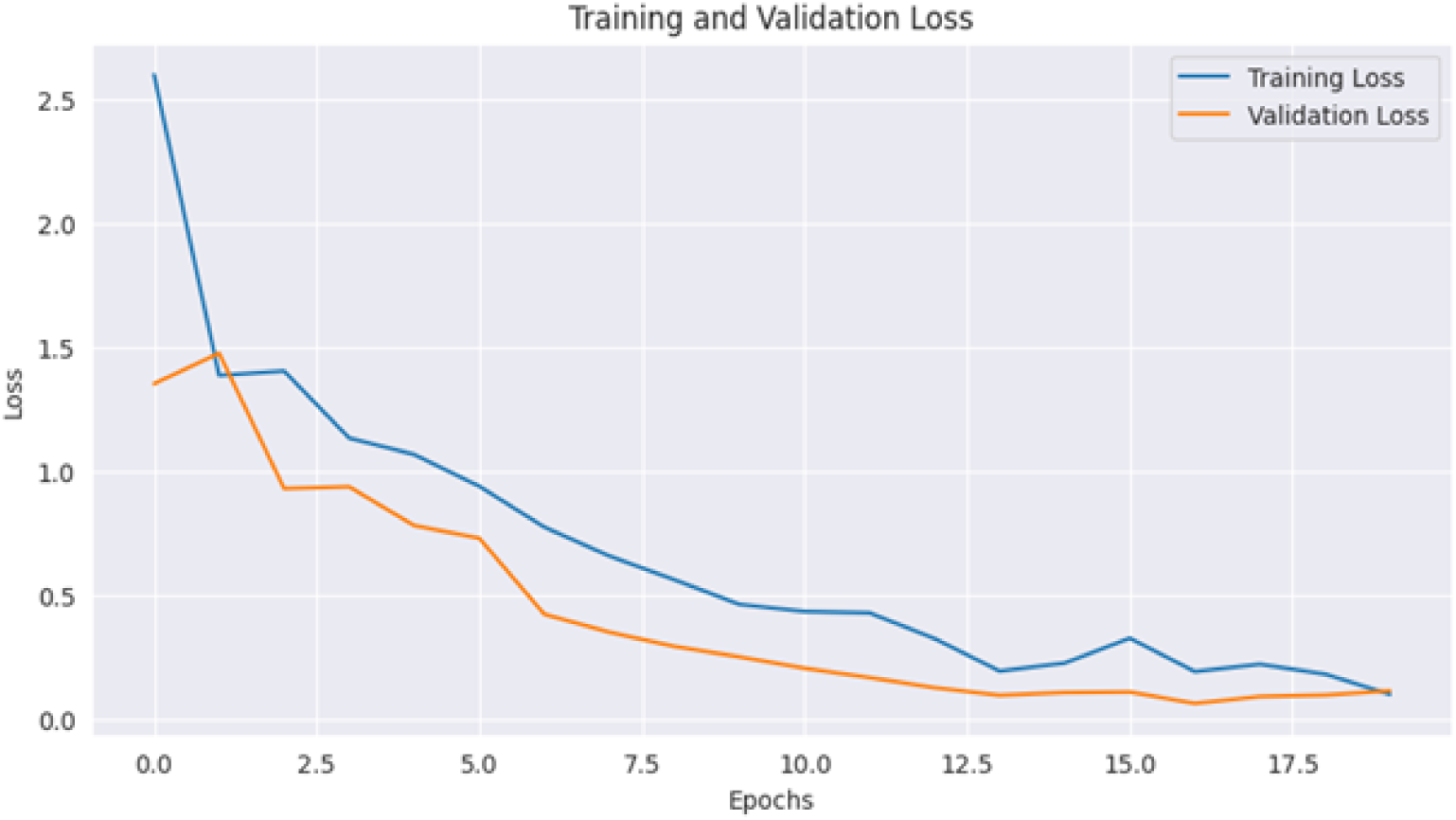
Training and validation loss over 20 epochs.

Convergence overall appeared smooth. Within the allocated training budget, performance reached a stable region, although minor validation fluctuations remained visible and were not analyzed further.

Since about 0.25 the training accuracy had improved and the ultimate epochs had an almost 0.96 accuracy. The accuracy of validation increased exponentially, and it did not decrease throughout the training. A nudging greater validation than training accuracy might seem counterintuitive in this arrangement, but nonetheless observed in practice. The process of training is augmented and regularized to introduce greater randomness, and validation is conducted more fixedly. This kind of tendency is common when the evaluation is intentionally harder than training is made to be [34].

In accordance with the trends of accuracy, the loss curves represent consistent optimization. The loss on training reduced to approximately 2.6 and validation loss reduced to approximately 0.1 and there are only minor peaks and valleys on the loss at subsequent epochs. No significant persistent generalization gap was observed and overfitting is thus said to be controlled as per the chosen training schedule and augmentation policy. Based on this, the validation macro-F1 is used in the selection of final checkpoints since multiple late checkpoints have the same level of performance, and the overall training dynamics are consistent.

### 0.13 Overall held-out test performance

Evaluation of the held out test set was done once after selecting the best checkpoint through validation macro-F1. The same preprocessing was used during training and testing, including resizing to 256 × 256 and ImageNet-style normalization, and the distribution of inputs was kept the same. The main report did not use any test-time augmentation or any calibration, so the metrics would reflect the raw classifier behavior. In the 15 labels, the macro-averaged performance is summarized in Table 5.

**Table 5.**
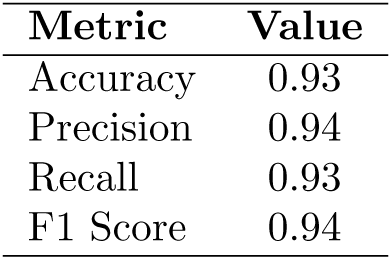
Overall held-out test set performance for the ResNet-50 + MSCA model (macro-averaged).

There is a high degree of proximity between the metric values implying fairly balanced behaviour among classes. Precision does not improve at the cost of recall nor do recall improvements incur at the cost of the precision. The fact that such balance exists implies that the remaining errors can be primarily due to the really hard visual cases, and not to the mere effect of thresholding or calibration, but this interpretation is not directly confirmed.

The results of the prediction on the held-out test set are summarized in Table 6. The overall accuracy of the test is 93.38% with 2,999 correct classifications, and 211 incorrect classifications. This summary offers a direct perspective of the absolute error volume in realistic field conditions, which adds to the reported macro-averaged metrics, but the behavior of the classes is not completely described.

**Table 6.** Overall prediction outcomes on the held-out test set.

| Prediction outcome | Number of samples |
| --- | --- |
| Correctly predicted samples | 2999 |
| Incorrectly predicted samples | 211 |
| Total test samples | 3210 |

### 0.14 The Controlled ablations: isolating MSCA, augmentation, and head effects

When clearly isolated as to the source of improvement, more confidence is gained as to the accuracy of the report. Controlled ablations were performed using the same data split, random seed, but with one factor held constant whilst the rest of the settings were held constant. This way, MSCA, augmentation, and classifier head design were analyzed separately and results of the test-sets were reported in Table 7. To maintain internal baseline the split, training schedule and evaluation protocol were kept constant and in this way, it was possible to attribute gains in performance more accurately than when comparing cross-papers though it is still required to assume that optimization behavior was maintained. Measurable gains are shown in Table 7 when MSCA is enabled under identical training conditions. With augmentation applied, macro-F1 rises from 0.920 to 0.937 (+0.017), while without augmentation an increase from 0.890 to 0.909 (+0.019) is observed. Under augmentation, a similar trend is followed by MCC, which increases from 0.912 to 0.929 (+0.017). Reported separately is an optional MixUp regularizer, whereas the default configuration—unless otherwise stated—combines MSCA with standard augmentation and a GAP-based head, a detail that is not always immediately clear from the table layout. Consistent improvement over the plain ResNet-50 baseline is therefore provided by MSCA when the protocol remains unchanged. Improved generalization under field variability is also associated with augmentation, and when augmentation is removed, a clear performance drop occurs for both the baseline and the MSCA-enhanced model. Third, final performance depends on the classifier head design, with GAP-based heads showing more stable behavior than flattening approaches, and the best overall variant under this protocol arising from a GAP-style configuration.

**Table 7.** Ablation study on the held-out test set using a fixed data split and controlled training settings.

| Setting | Acc. | Macro-P | Macro-R | Macro-F1 | MCC |
| --- | --- | --- | --- | --- | --- |
| ResNet-50 (no MSCA) + augmentation | 0.919 | 0.923 | 0.918 | 0.920 | 0.912 |
| ResNet-50 + MSCA + augmentation | **0.934** | **0.941** | **0.933** | **0.937** | **0.929** |
| ResNet-50 (no MSCA), no augmentation | 0.892 | 0.895 | 0.889 | 0.890 | 0.879 |
| ResNet-50 + MSCA, no augmentation | 0.908 | 0.912 | 0.906 | 0.909 | 0.898 |
| Head: Flatten (MSCA + augmentation) | 0.925 | 0.931 | 0.924 | 0.928 | 0.919 |
| Head: GAP (MSCA + augmentation) | 0.930 | 0.936 | 0.929 | 0.932 | 0.924 |

### 0.15 Multi-seed evaluation and result stability

Stability beyond a single initialization was assessed utilizing a multi-seed evaluation. The same fixed train–validation–test split was retained, and the experiment was repeated with three different random seeds. The data split was unchanged, similarly, the Preprocessing and augmentation were kept the same. Optimizer settings and the learning schedule were also kept unchanged. This design isolates the effect of random initialization. Stable performance across different random initializations is indicated by the small standard deviations reported in Table 8. It is therefore suggested that the observed gains are not driven by a single favorable seed, but instead reflect consistent improvements linked to the architectural design, although other sources of variance were not examined explicitly.

**Table 8.** Multi-seed evaluation results (mean ± standard deviation) over three independent runs using the same fixed data split.

| Metric | Mean | Std. |
| --- | --- | --- |
| Accuracy | 0.934 | 0.004 |
| Macro-Precision | 0.941 | 0.003 |
| Macro-Recall | 0.933 | 0.004 |
| Macro-F1 | 0.937 | 0.003 |

**Table 9.** Statistical significance testing using paired t-test across three independent runs.

| Comparison | Metric | p-value |
| --- | --- | --- |
| ResNet-50 vs. ResNet-50 + MSCA | Macro-F1 | < 0.05 |

### 0.16 Statistical significance analysis

To determine whether the model behavior was consistent as opposed to stochastic variation, statistical significance testing was conducted in order to ascertain whether the observed improvements were a reflection of the model behavior or not. The reason paired testing was considered suitable to this environment was that all the variants were trained and tested on the same fixed data split. A paired t-test was used to compare the ResNet-50 +MSCA with the baseline of ResNet-50 no MSCA module. The data were in the form of Macro-F1 scores that were obtained after three separate runs, with each run being initialized in a different random seed, and the training and evaluation settings being the same. The importance of the different performances in the null hypothesis of same model performance is indicated by the p -values obtained, but the number of runs is very low and may limit the statistical power of the analysis. The statistical analysis supported the fact that the performance increase achieved when the MSCA module gets implemented is statistically significant at the 95% confidence level. This is consistent with the conclusion that the apparent gains are due to the architecture design and not the random variation between the different runs; however, better accuracy of the estimate can be achieved by adding more random seeds.

### 0.17 Comparison with established deep learning baselines

The effectiveness of the MSCA enhanced architecture was validated by carrying out direct comparative analyses. The assessments were conducted on EfficientNet and a Vision Transformer, with attention based convolutional neural network being used as an extra baseline. Training was done on all the models with the same fixed data division, the same preprocessing, augmentation, and optimizer. The evaluation protocol was applied equally to each model to identify comparability and also to counter the confounding variability, but still, there can be residual differences in optimization behaviors that can be obtained since there was architectural variance. Under field imagery circumstances of clutter, varying illumination and a mix of scale, recognition performance was measured therefore permitting the variations to be ascribed more to architectural design and not necessarily to variations in the training arrangement. As shown in Table 10 the proposed ResNet-50 + MSCA model is outperforming the evaluated baselines across the reported metrics. The gain is being observed most clearly in macro-F1, reflecting balanced behavior across classes, with varying visual difficulty. Transformer-based models are providing strong representational capacity, but their performance, under the given data regime and field variability, is remaining slightly lower than the proposed approach. Overall, the results are suggesting that combining explicit multi-scale contextual aggregation with lightweight attention refinement is remaining effective, particularly for fine-grained agricultural image classification, when complex and cluttered scenes are being handled.

**Table 10.** Quantitative comparison with established deep learning baselines on the held-out test set. All models were evaluated under the same training and evaluation protocol.

| Model | Accuracy | Macro-P | Macro-R | Macro-F1 |
| --- | --- | --- | --- | --- |
| EfficientNet-B0 | 0.912 | 0.916 | 0.910 | 0.913 |
| EfficientNet-B3 | 0.918 | 0.922 | 0.917 | 0.919 |
| Vision Transformer (ViT-B/16) | 0.905 | 0.909 | 0.902 | 0.905 |
| Attention-based CNN (CBAM) | 0.921 | 0.926 | 0.920 | 0.923 |
| **Proposed ResNet-50 + MSCA** | **0.934** | **0.941** | **0.933** | **0.937** |

### 0.18 Per-class behavior and error patterns on a balanced diagnostic subset

Aggregate metrics can hide which classes are difficult and why they fail. To make per-class precision and recall directly comparable, we evaluate on a balanced diagnostic subset containing 50 images per class, for a total of 750 images. This diagnostic subset supports clearer error interpretation because differences are not confounded by class frequency. The Table 11 reports per-class precision, recall, F1, and support scores.

**Table 11.**
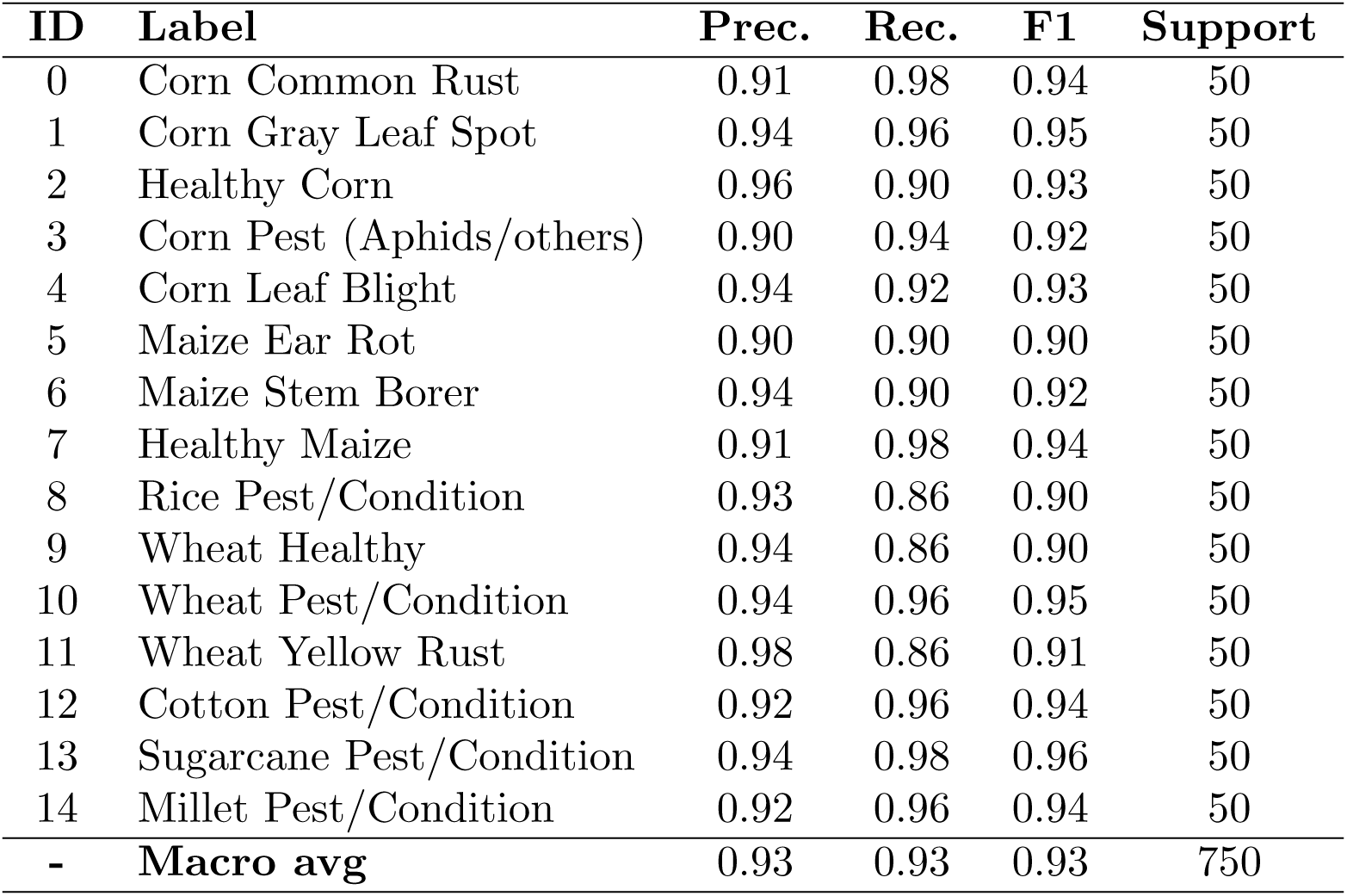
Per-class precision/recall/F1 on a balanced diagnostic subset (50 images per class; total 750).

Some of the classes have very high recall with Corn Common Rust, Healthy Maize, and Sugarcane Pest/Condition recording 0.98 recall with high F1s. In the case of these labels, performance is constantly high. Much more difficult behavior is observed in Rice Pest/Condition, Wheat Healthy and Wheat Yellow Rust whereby recall is 0.86 and the precision is high, indicating a conservative decision behavior. The pattern can be attributed to field situations where the symptoms are low, spatially small, or even overlapping in a visual image with different illumination. The noted error patterns have typical features of field imagery. The lesions that occupy a small fraction of the image are commonly missed, lesions that overlap foliage partially are sometimes missed, and lesions that are underbacklit are sometimes missed because of low contrast. In such cases, attention can occasionally disperse in other parts of the leaf such as veins and areas of homogenous texture as opposed to highlighting edges of lesions, a factor that complicates the separation of homogenous conditions. Specific augmentation of small or low-contrast symptoms is thus driven, along with more-resolution or tiling inference in case deployment supports that. Attention-based visualizations were subject to qualitative analysis to determine model focus. The annotations of lesions at pixel level were not provided and quantitative analysis was not performed because expert validation was not performed. Nevertheless, the focus is still more likely to be around lesion areas or pest clusters in correctly rated samples, and diffuse attention across veins or smooth areas is found in false ones. The results of such observations can serve as qualitative data to prove that MSCA contributes to symptom-oriented learning, but there are still cases when attention refinement is quite difficult.

### 0.19 Effect of backbone depth under a controlled setting

Recognition performance is influenced by backbone depth, particularly in field imagery where subtle textures and strong scale variation are present [35]. To quantify this effect, three residual backbones were trained using the same data split, augmentation policy, loss function, and early-stopping criterion, while MSCA was kept enabled throughout. Variation was restricted to backbone depth only. Reported in Table 12 are the held-out accuracy and loss for ResNet-18, ResNet-34, and ResNet-50, enabling a controlled comparison that isolates depth-related effects. The highest accuracy and the lowest loss are achieved by ResNet-50 in this controlled comparison. Likely captured by the deeper backbone are stronger hierarchical features, which becomes relevant when overlapping textures and small symptom regions are common in field imagery. Although a lower loss is observed for ResNet-34 than for ResNet-18, its accuracy is slightly reduced, a behavior that may arise from differences in confidence calibration under cross-entropy optimization. Based on these observations, ResNet-50 is retained as the default backbone for the proposed MSCA-enhanced recognition system.

**Table 12.** Held-out test accuracy and loss for residual backbones (all with MSCA enabled).

| Model | Accuracy (%) | Loss |
| --- | --- | --- |
| ResNet-18 | 90.71 | 0.1610 |
| ResNet-34 | 89.42 | 0.1119 |
| ResNet-50 | 93.38 | 0.06153 |

**Table 13.**
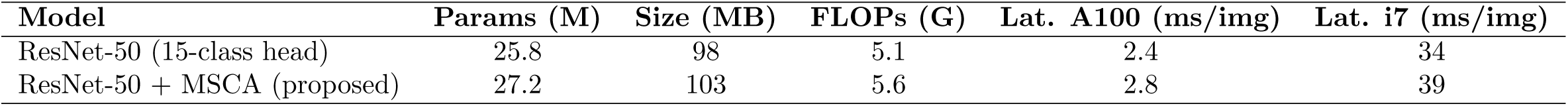
Deployability-oriented profiling at 256 × 256 input. FLOPs are estimated; latency is the median forward-pass time after warm-up.

### 0.20 Deployment-oriented profiling

The usability in the context of real-world precision-agriculture processes cannot be guaranteed only with the high accuracy; there are other limitations like manageable memory usage, affordable computational complexity, and the latency that makes timely decisions, among others, which need to be met in practice. With the configuration as seen in Table 4, the number of parameters, the size of the approximate model, the number of FLOPs, and the median inference time at the resolution of 256 × 256are as follows. Mean forward pass latency is the forward pass latency at the end of the warm-up period, and omit any data loading and preprocessing delays, but remain potentially significant to the overall end to end timing in real system implementations. The MSCA presents a quantifiable change in the computational load, but the deployment requirements are met. Since inference is possible on smartphones, drones, or any other embedded agricultural systems, efficiency is a major priority. Modern GPUs can preserve near-real time performance with the extra attention branches, and can be deployed to the edge with feasibility through optimization. The moderate increase in the count of parameters and FLOPs points to the trade-off between contextual refinement and efficiency which is considered to be acceptable within the fixed evaluation protocol. Parallel convolutional filters and lightweight attention projections are added leading to slight growth in the number of parameters. FLOPs increase as well due to enlargement of receptive-fields by dilated branches; however, at the chosen 256 by 256 resolution, still, the expansion is restrained. On a GPU, A100, it achieves a low per-image latency; on an i7 CPU it exhibits higher latency, but appreciable latency reduction can be achieved by use of quantization, FP16 inference, or run-time optimization. Even though the results of profiling vary depending on the specifics of implementation, they show that, under the right deployment settings, it is possible to support near-real-time field monitoring..

### 0.21 Indicative comparison with recent literature

The beneath Table 14 summarizes some of the recent research results which are simply given to put the research findings in context. This comparison is not supposed to be a strict standard, but rather an indication one, as datasets, label spaces, acquisition conditions and train test protocols differ significantly among existing literature. Direct numerical comparison is not necessarily always significant especially since most studies use laboratory-type datasets or smaller sets of classes. Instead of arguing that it is absolutely better than a heterogeneous evaluation environment, performance on field-style imagery with a richer label taxonomy is put in context. Many existing methods have been used to report results on controlled datasets with cleaner backgrounds and lower levels of environmental variability. Contrarily, the suggested framework is tested on field imagery where the background is cluttered, there are partial occlusions, and sudden illumination variations, with covering a larger crop and pest/condition label space. In such practical circumstances, the achieved accuracy measures the strength of the learnt feature representations and not performance in the case of idealized acquisition conditions, but the comparison is not necessarily coincidental. Under field-style acquisition and a 15-class taxonomy, 93.38% accuracy is achieved by the proposed approach. Effective discrimination of visually similar conditions in complex scenes is enabled by combining a residual backbone with multi-scale contextual attention. Caution, however, should be exercised when interpreting cross-study comparisons, since variations in label definitions and experimental protocols can shift reported metrics. Unlike aggregated cross-paper summaries, direct and controlled baselines are provided in Section 0.17, where modern CNN and transformer-based architectures are evaluated under identical conditions.

**Table 14.** Representative plant/crop recognition results in the literature (indicative comparison).

| Reference | Classes | Dataset source | Conditions | Model | Accuracy (%) |
| --- | --- | --- | --- | --- | --- |
| [7] | 4 | Own collected | In-field | Inception/VGG variant | 80.38 |
| [8] | 4 | PlantVillage | Lab | VGG + Inception modules | 84.25 |
| [6] | 4 | PlantVillage | Lab | CNN (author design) | 90.80 |
| [5] | 8 | Own collected | In-field | Custom CNN | 92.85 |
| **Ours** | 15 | Curated dataset | In-field | **ResNet-50 + MSCA** | **93.38** |

### 0.22 Summary, limitations, and practical implications

Overall, the main design choice is supported by the reported results. Under a controlled protocol, consistent improvements are yielded by MSCA over the non-attention baseline, while robust invariances under field variability are actively learned through augmentation. Where performance is weaker is clarified by the class-wise diagnostic analysis, with recall reductions being concentrated in cases involving small or low-contrast symptoms. In unconstrained field imagery, such failure modes are expected. Several limitations are also acknowledged. From a single curated source, all images were obtained, and cross-dataset or cross-region evaluation was not performed; therefore, performance under unseen geographic regions, imaging devices, or seasonal variation cannot be directly quantified. As image-level classification, the task is formulated, and pixel-level or region-level annotations are not available, which limits quantitative evaluation of localization accuracy and attention alignment. Across pest or condition categories, some labels were aggregated; granularity was reduced by this choice, but sufficient sample sizes and annotation reliability were preserved. The findings are not invalidated by these constraints, although directions for future work are clearly motivated. An important deployment concern remains generalization beyond the evaluated dataset. Substantial variation is exhibited by field imagery in lighting, background clutter, occlusion patterns, crop varieties, and acquisition devices. Although diverse field-style images are included, explicit external evaluation was not conducted. The reported results should therefore be interpreted as evidence of robustness within the current data domain rather than as a guarantee of cross-domain transfer, and future work should incorporate cross-dataset and cross-season validation to assess transferability more directly. Generalization is still targeted by the adopted design choices. Field-style imagery with strong variability is used during training, while augmentation is applied only to the training subset. Contextual and scale-robust features are emphasized by attention refinement, instead of learning dataset-specific shortcuts. Transfer to related agricultural monitoring scenarios is expected to be supported by these factors, although confirmation through external validation remains necessary. Practical improvements are also suggested by the observations. Tiny lesions and low-contrast patterns can be amplified through targeted augmentations. For early-stage symptoms, higher-resolution or tiled inference may be employed when deployment budgets allow. Quantitative interpretability evaluation would be enabled by region-level annotations. With such refinements, improved reliability can be achieved by the unified pipeline for real monitoring workflows, while reproducibility and deployment-oriented evaluation are preserved.

## Conclusion

An image-based system for unified crop and pest/condition recognition was evaluated under realistic field conditions, using a ResNet-50 backbone augmented with a lightweight Multi-Scale Contextual Attention (MSCA) module and trained on 21,404 images across 15 classes with a fixed, leakage-aware protocol. During training, stable convergence was observed, and clarity was improved through more explicit description of the MSCA design, dataset preparation, and evaluation strategy. On the held-out test set, an accuracy of 0.93 and a macro-F1 of 0.94 were achieved, with precision and recall remaining closely aligned. Under identical settings, controlled ablations demonstrated consistent gains over the non-attention baseline, and from the backbone-depth comparison it was evident that ResNet-50 outperformed shallower variants. Class-wise behavior was largely even. Lower recall nevertheless appeared in a small number of labels involving tiny, faint, or occluded symptoms, where attention sometimes diffused toward veins or uniform texture regions rather than focusing sharply. Suitable for decision support in field scouting is the overall pipeline, which also compares favorably with laboratory-focused studies. Still, it remains limited by the absence of cross-dataset evaluation and explicit localization, motivating future work on targeted augmentation, higher-resolution inference, and broader regional validation.

